# DNA methylation is a widespread mechanism of light-induced circadian clock period plasticity

**DOI:** 10.1101/2022.06.30.498269

**Authors:** Suil Kim, Douglas G. McMahon

## Abstract

The suprachiasmatic nucleus (SCN) of the hypothalamus is a principal light-responsive circadian clock that adjusts circadian rhythms in mammalian physiology and behavior to changes in external light signals. Although mechanisms underlying how light acutely resets the timing of circadian rhythms have been characterized, it remains elusive how light signals induce lasting changes in circadian period, so-called period after-effects. Here we have found that the period after-effects on circadian behavior of changing photoperiods are blocked by application of DNA methyltransferase inhibitors directed to the SCN. At the level of single light pulses that act as clock-resetting stimulations, pharmacologically inhibiting DNA methylation in the SCN significantly attenuates period after-effects following acute phase shifts in behavioral rhythms in vivo, and blocks period after-effects on clock gene rhythms in the isolated ex vivo SCN. Acute clock resetting shifts themselves, however, do not appear to require DNA methylation at the SCN and behavioral levels, in contrast to subsequent period plasticity. Our results indicate that DNA methylation in the SCN mediates light-induced period after-effects in response to photoperiods, and single light pulses, and together with previous studies showing that DNA methylation in the SCN is essential for period after-effects of non-24hr light cycles (T-cycles), suggest that DNA methylation in the SCN is a widespread mechanism of light-induced circadian period plasticity.

## Introduction

Circadian rhythms are pervasive in mammalian physiology and behavior in anticipation of, and alignment with, daily environmental cycles. Nearly all mammalian cells including neurons and glia have an endogenous 24-hour timing mechanism, or circadian clock, the molecular basis of which is self-sustained circadian oscillations of clock genes including *Period* (*Per*) and *Cryptochrome* (*Cry*) genes via autoregulatory transcription-translation feedback mechanisms (Takahashi, 2017). Individual cellular and tissue circadian rhythms in the body are adjusted in tune with daily light cycles by the suprachiasmatic nucleus (SCN) in the hypothalamus which receives retinal light input(Welsh et al., 2010). The period length of circadian rhythms, or circadian period, is species-specific and genetically defined, but can be enduringly modified by the effects of external light signals (Pittendrigh and Daan, 1976a, 1976b). Light exposures at night which cause acute delays or advances in the timing (phase) of circadian rhythms also subsequently lead to enduring changes in circadian period, or so-called period after-effects. Light-induced phase delays cause period lengthening after-effects, while phase advances cause period shortening after-effects (Pittendrigh and Daan, 1976a). In addition, synchronization or entrainment of circadian rhythms to different light-dark (LD) cycles, such as altered durations of daylight (photoperiods; e.g., 16 hr light and 8 hr dark per 24 hr [LD 16:8]), and non-24h light-dark cycles (T-cycles; e.g., 11 hr light and 11 hr dark per 22 hr [T22]), cause pronounced period after-effects (Pittendrigh and Daan, 1976b).

Previous work (Azzi et al., 2014) showed that DNA methylation in the mouse SCN is necessary for expression of after-effects on circadian period of behavioral rhythms following weeks of entrainment to non-24h light cycles, suggesting that epigenetic regulation of gene expression might mediate long-lasting changes in circadian clock properties. However, it remains an open question whether DNA methylation is involved in period after-effects that arise from the photoperiodic variation in 24-hour light cycles that is experienced by many organisms and people across the seasons. Also, it remains unclear whether DNA methylation regulates period after-effects at the level of single light pulses or clock-resetting cues, or it does so only after chronic light exposures such as light entrainment.

Here, we show that pharmacological inhibition of DNA methylation in vivo directed near the SCN significantly attenuates photoperiodic aftereffects on the circadian period of behavioral rhythms, and uncover that DNA methylation regulates period plasticity of SCN clock gene rhythms and behavioral output rhythms at the level of acute clock-resetting stimulations without influencing induced phase shifts. Our results suggest that DNA methylation is a fundamental mechanism of circadian plasticity to prior light history.

## Results

### Inhibition of DNA methylation targeted to the SCN blunts the after-effects of photoperiods on circadian behavior

Entrainment of mouse circadian locomotor behavior to long summer-like photoperiods (e.g., 16 hr light per 24 hr) causes a subsequent shortening of the endogenous circadian period when measured in constant darkness (Pittendrigh and Daan, 1976b). To test whether DNA methylation is involved in period after-effects of long photoperiod entrainment, we infused RG108, a pan-inhibitor of DNA methyltransferases, into the third ventricle adjacent to the SCN during entrainment of wheel-running behavior rhythms to LD 16:8 photoperiods (Figure 1A). This method of infusion was previously shown to effectively target the SCN (Azzi et al., 2014). Mice which had been maintained in LD 12:12 cycles were placed in running wheel cages and allowed to free-run for 7 days in constant darkness to establish their baseline circadian locomotor rhythm period. Mice were then re-entrained to LD 12:12 cycles for 9-10 days, with infusion of RG108 started on day 4 or 5, and either continued for an additional 10 days in LD 12:12 (Control), or entrained to LD 16:8 cycles for 12 days (LD 16:8), and then allowed to free-run for 7 days to assay for period after-effects. Control group RG108-infused mice did not show significant differences in endogenous period change compared to vehicle controls (Figure 1B), suggesting that inhibition of DNA methylation does not itself affect the baseline circadian period. In contrast, LD 16:8 vehicle control mice exhibited the expected shortening of their circadian locomotor period, which was significantly blunted by RG108 (Figure 1), suggesting that DNA methylation regulates changes to circadian period following photoperiodic entrainment.

**Figure 1.**
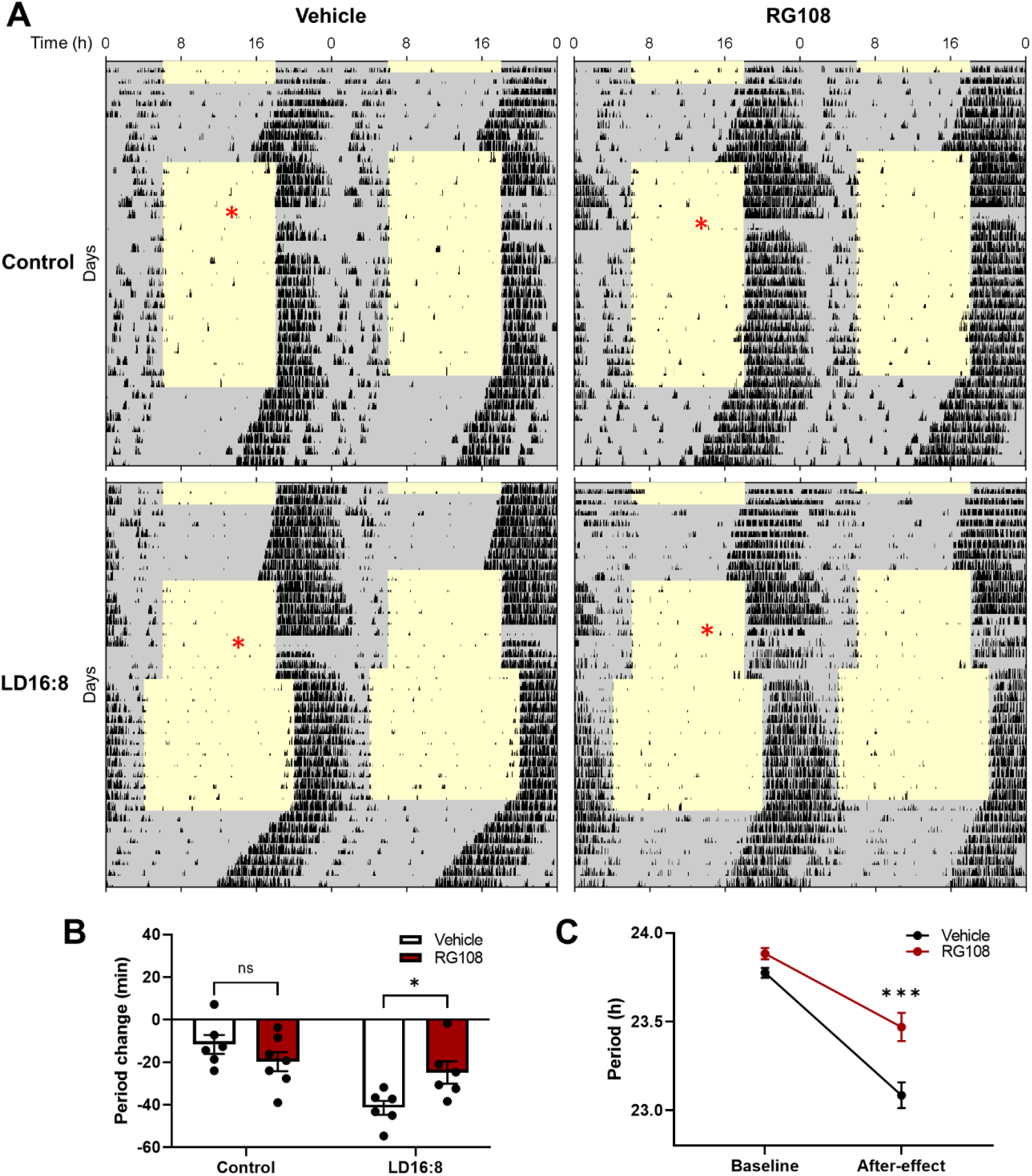
DNA methylation inhibitor attenuates after-effects on circadian period following long photoperiod entrainment. **(A)** Representative double-plotted mouse locomotor actograms under different lighting conditions (LD 12:12 cycle, LD 16:8 cycle, constant darkness). Yellow and grey areas indicate light and dark phases, respectively. Black ticks indicate wheel-running behavior patterns. Red asterisks denote onset of RG108 or vehicle infusion into the third ventricle of the brain. **(B)** Quantification of changes in the endogenous period in constant darkness in control (LD 12:12 cycle) and LD 16:8 (change from LD 12:12 to LD 16:8 cycle) conditions with RG108 or vehicle infusion. (Two-way ANOVA with Sidak’s multiple comparisons tests, mean ± SEM, n = 6-7, ns: not significant, *p < 0.05). **(C)** Quantification of the endogenous period at baseline (the first constant darkness) and at the after-effect period (the second constant darkness) following long photoperiod entrainment in the presence of RG108 or vehicle infusion. (RM two-way ANOVA with Sidak’s multiple comparisons tests, mean ± SEM, n = 6, ***p<0.001).

### Expression of period after-effects in circadian behavior following acute phase shifts depends on DNA methylation

Resetting of circadian rhythms by discrete light exposure is fundamental for circadian entrainment to light cycles (Pittendrigh and Daan, 1976b). In addition, acute phase delays or advances in circadian rhythms induced by brief light exposures cause period after-effects, with period lengthening or shortening on subsequent cycles (Pittendrigh and Daan, 1976a). Interestingly, the magnitude of subsequent period changes is positively correlated with the magnitude of the preceding phase shifts (Sharma and Daan, 2002; Sharma, 2003). We thus sought to investigate two questions. First, is DNA methylation involved in period changes following acute light exposure, not just following weeks of entrainment to different photoperiods (Figure 1), or non-24h light cycles (Azzi et al., 2014)? Second, does DNA methylation regulate period after-effects by modulating the magnitude of initial phase shifts, or rather by directly acting on expression of period after-effects?

To test whether and how DNA methylation mediates period after-effects at the level of discrete light pulses, we sought to inhibit DNA methylation in the mouse brain with RG108 injection into the third ventricle near the SCN, and then delivered 1h light pulses in the early physiological night (∼circadian time (CT) 13.5, ∼CT16.5) in constant darkness to induce phase shifts and period after-effects in locomotor behavior rhythms (Figure 2A). Vehicle control mice showed phase delays and period lengthening following light pulses, as expected (Figure 2B-D). RG108-treated mice showed similar magnitude phase delays to the vehicle controls, but period lengthening was blocked (Figure 2B-D), indicating that DNA methylation regulates period changes without directly affecting phase shifts. This supports an idea that DNA methylation mediates expression of period after-effects following acute resetting of circadian rhythms.

**Figure 2.**
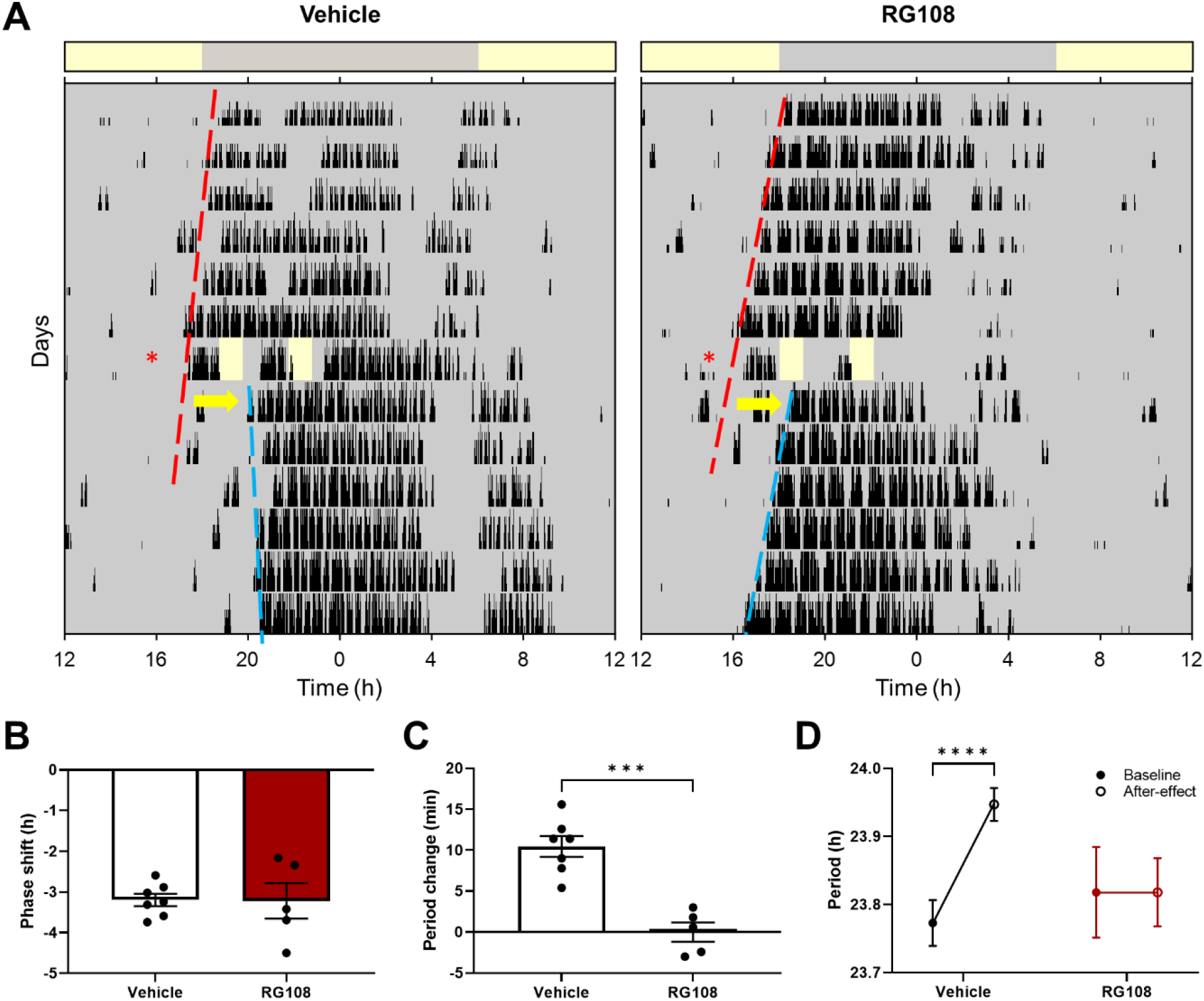
DNA methylation mediates after-effects on circadian period of locomotor behavioral rhythms without affecting acute phase delays. **(A)** Representative wheel-running behavior actograms with brief light exposure (yellow bars) in constant darkness with a drug vehicle (left) or RG108 (right) injection (red asterisk) in the third ventricle. Bars above the actograms indicate the 12h light (yellow) and 12h dark (grey) cycle previously entrained to which mice were before release into constant darkness. Red and blue dashed lines indicate linear regression of locomotor activity onsets before and after light exposure, respectively. The slopes of regression lines are the calculated activity rhythm periods. Yellow arrows denote phase shifts. **(B-C)** Quantification of phase shifts **(B)** and period changes **(C)** following light exposure in vehicle- and RG108-injected mice. (Unpaired t-test, mean ± SEM, n = 5-7, ***p<0.001). **(D)** Quantification of the endogenous period before and after light exposure (baseline, after-effect) in vehicle- and RG108-injected mice. (RM two-way ANOVA with Sidak’s multiple comparisons tests, mean ± SEM, n=5-7, ****p<0.0001).

### After-effects of phase resetting on the period of SCN molecular rhythms are modulated by DNA methylation

The preceding experiments in intact animals showed that inhibition of DNA methylation directed at the SCN blocks expression of behavioral period plasticity to photoperiodic entrainment and to single phase shifts. As circadian behavioral rhythms are regulated by interactions between the central SCN clock and other peripheral brain clocks (Kalsbeek et al., 2006; Begemann et al., 2020), in principle, DNA methylation might be acting directly on expression of period plasticity in the SCN itself, or through actions on other brain clocks downstream of the SCN for behavioral outputs.

To test whether DNA methylation mediates circadian after-effects directly on SCN molecular rhythms, we used organotypic SCN slice cultures expressing a bioluminescent reporter of clock protein PER2 expression (PER2::LUC; Yoo et al., 2004), and induced phase shifts in PER2::LUC rhythms in SCN slices by applying vasoactive intestinal peptide (VIP), a neuropeptide mediating light-induced SCN clock resetting(Welsh et al., 2010), at early physiological night (CT14) in the presence or absence of the DNA methylation blocker RG108 in the culture medium (Figure 3). RG108 treatment itself without VIP application did not change the phase or period of PER2::LUC bioluminescence rhythms in SCN slices (Figure 3A-C), suggesting that DNA methylation is not essential for maintaining SCN rhythms in the resting state. This is consistent with previous observations (Azzi et al., 2014) and our results here that blocking DNA methylation itself does not affect the endogenous period of circadian behavior. When VIP was applied to SCN slices, RG108 co-treatment did not affect VIP-induced phase shifts in SCN rhythms, but it attenuated VIP-induced period changes (Figure 3A-C). This suggests that DNA methylation directly modulates expression of SCN rhythm after-effects following acute phase shifts, consistent with our findings in behavioral experiments.

**Figure 3.**
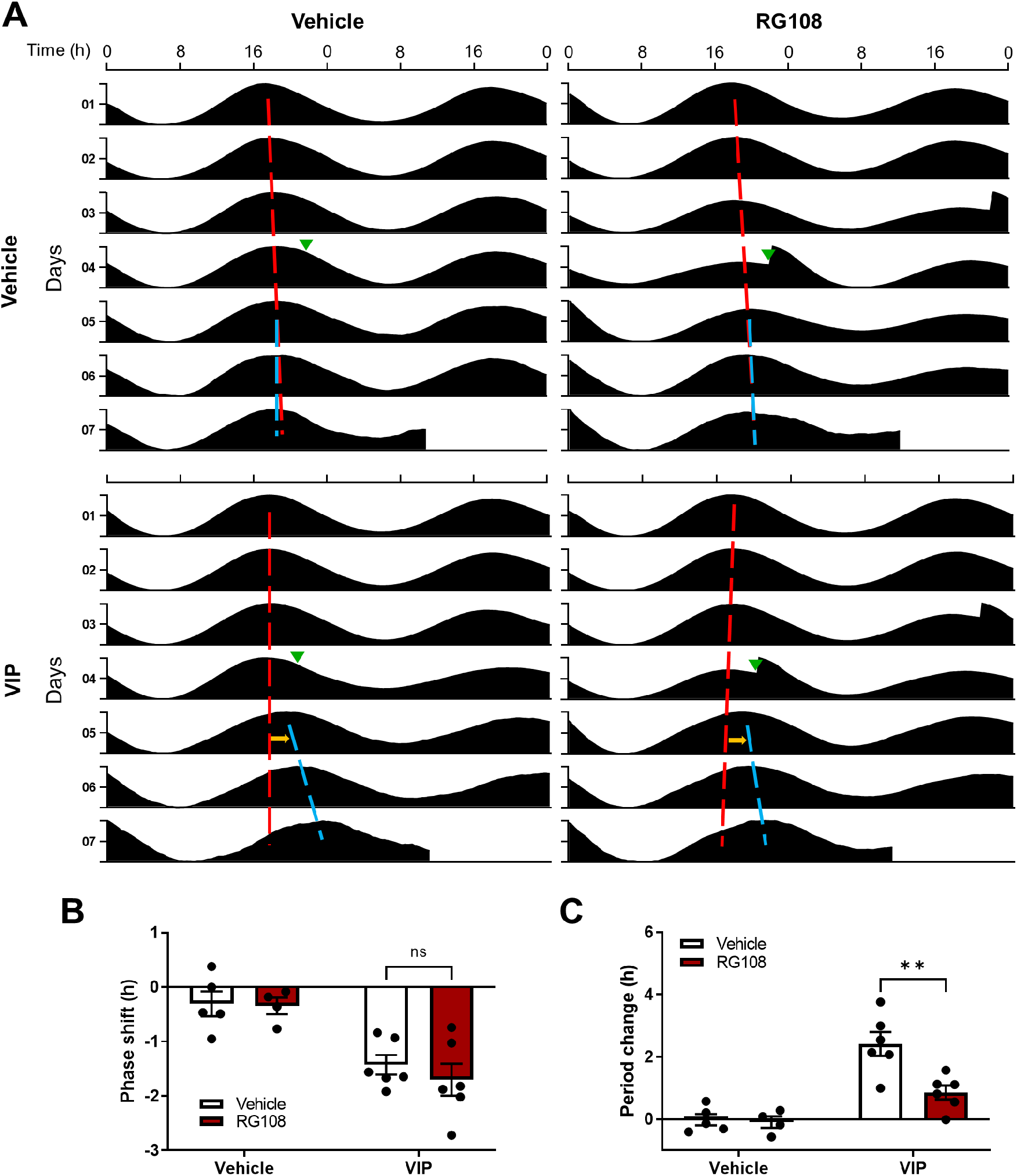
DNA methylation mediates VIP-induced after-effects on the ex vivo SCN rhythm period without affecting acute phase delays. **(A)** Representative double-plotted actograms of PER2::LUC bioluminescence rhythms in SCN slices treated with one of combinations of VIP, RG108, and their vehicles. Red and blue lines indicate linear regression of peaks before and after treatment (green triangles), respectively. Yellow arrows denote phase shifts. **(B-C)** Quantification of phase shifts **(B)** and period changes **(C)** following treatment. (Two-way ANOVA with Sidak’s multiple comparisons tests, mean ± SEM, n=4-6, ns: not significant, **p<0.01).

Since in the previous experiments VIP was applied as a bolus and not washed out, its continued presence in the medium could potentially confound after-effects on period with persistent ongoing effects of VIP itself on period. Therefore, we sought to test these results with a more temporally precise stimulus to the ex vivo SCN. Recently, our lab showed that optogenetic stimulation of the ex vivo SCN using a red light-sensitive opsin ChrimsonR mimics light-induced resetting of circadian rhythms in vivo (Kim and McMahon, 2021). Here, we expressed ChrimsonR in neurons in SCN slices from PER2::LUC mice with an AAV (AAV-Synapsin-ChrimsonR-tdTomato; Klapoetke et al., 2014). We then applied to the SCN slices a different class of pan-DNA methyltransferase inhibitor, SGI-1027, and one day later delivered 15-minutes of optogenetic stimulation (625 nm, 10 Hz, 10 ms pulse width) to SCN neurons in their early physiological night (CT14) to mimic retinal light input to the SCN (Figure 4). SGI-1027 treatment itself without optogenetic stimulation did not affect SCN PER2::LUC rhythms (Figure 4A-C), suggesting again that DNA methylation does not influence resting-state circadian rhythms in the SCN. SGI-1027 treatment prior to optogenetic stimulation suppressed period changes in the SCN rhythms following the stimulation, but it did not impact phase shifts (Figure 4A-C). This further supports the notion that DNA methylation is not essential for phase shifts in SCN rhythms following acute light input, but it is required for expression of after-effects on SCN rhythm period on subsequent cycles.

**Figure 4.**
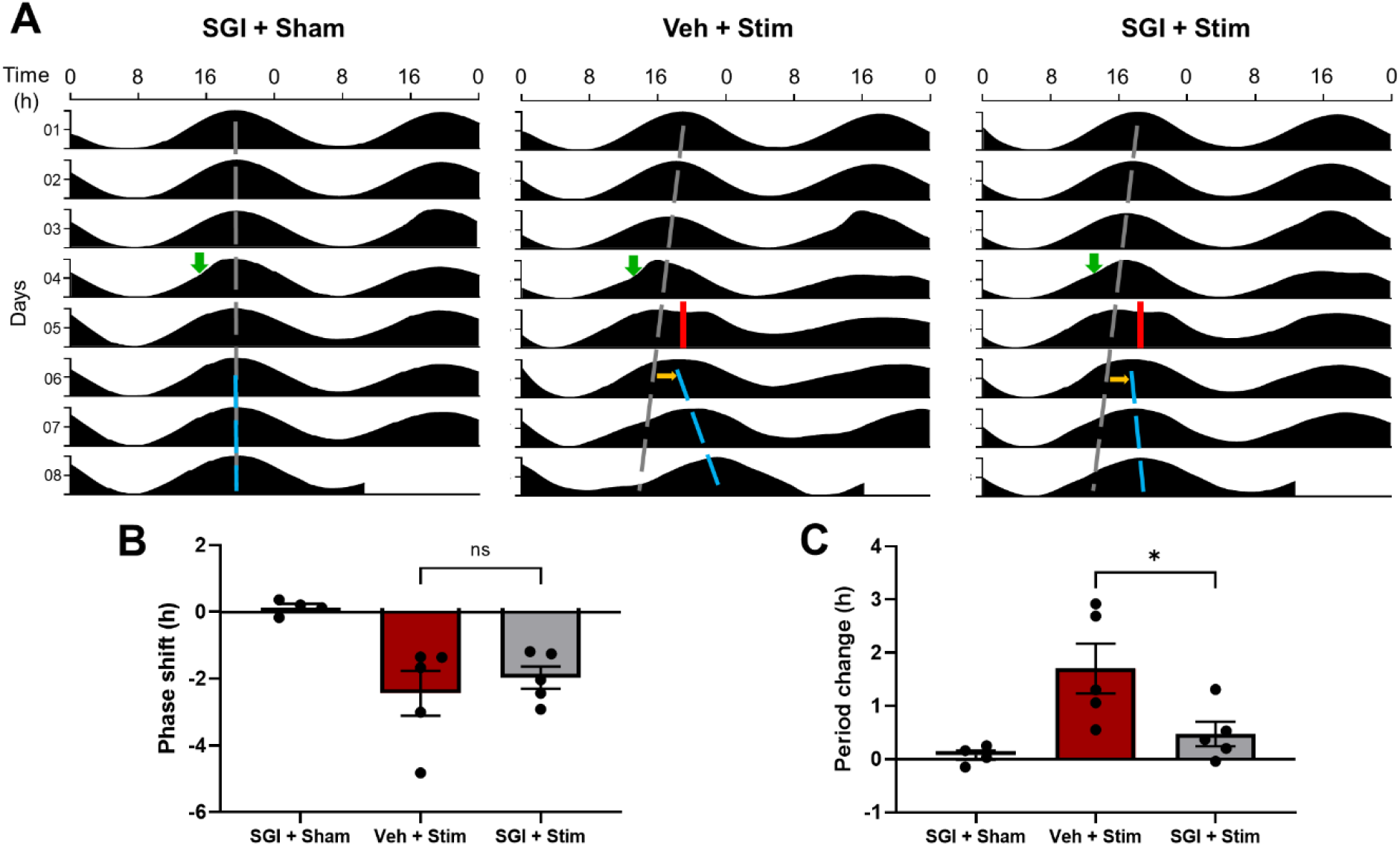
DNA methylation is critical for after-effects on the ex vivo SCN rhythm period following acute optogenetic stimulation of SCN neurons. **(A)** Representative double-plotted actograms of PER2::LUC bioluminescence rhythms in SCN slices with SGI-1027 or its vehicle treatment (green arrows), and sham or optogenetic stimulation (red bars). Grey and blue dashed lines indicate linear regression of peaks before and after stimulation, respectively. Yellow arrows denote phase shifts. **(B-C)** Quantification of phase shifts and period changes **(C)** following sham or optogenetic stimulation. (One-way ANOVA with Tukey’s multiple comparisons tests, mean ± SEM, n=4-5, ns: not significant, *p<0.01).

## Discussion

This study extends the understanding that DNA methylation is involved in light modulation of circadian period, a lasting form of plasticity in the mammalian brain biological clock. Here we have shown that entrainment to altered daytime lengths (photoperiods) involves DNA methylation to express after-effects on circadian period, and uncovered that at the level of single light pulses the SCN expresses circadian after-effects via DNA methylation. Thus, our results suggest that the SCN involves DNA methylation as a widespread mechanism of period after-effects at the molecular and behavioral output levels.

After-effects on circadian period is a prominent example of circadian clock plasticity to different lighting conditions. DNA methylation was previously shown to be necessary for period plasticity in the specific case of T-cycle entrainment (Azzi et al., 2014, 2017). However, the role of DNA methylation in period plasticity following other forms of light exposure remained uncharacterized. Unlike T-cycle entrainment, which is rarely, if ever, experienced in nature, photoperiod entrainment is a key function of the circadian clock critical for mediating seasonal changes in animal physiology and behavior. We now show that period plasticity following photoperiod entrainment likely involves DNA methylation in the SCN. Interestingly, DNA methylation plays a key role in regulating seasonal timing of reproduction in many different taxa including animals and plants (Viitaniemi et al., 2019). In rodents, DNA methylation is critical for hypothalamic regulation of seasonal reproductive organ development (Stevenson and Prendergast, 2013). This suggests that seasonal changes in DNA methylation patterns in different brain regions, including the SCN, might coordinately regulate seasonality of animal physiology and behavior. Future studies will be needed to investigate how photoperiod-induced period plasticity is coordinated with other hypothalamic regulation for seasonal physiology and behavior.

Another important point of this study is that our data suggest that DNA methylation can take place in the SCN following even a brief light exposure or a clock-resetting stimulation to mediate period plasticity. Given that DNA methylation is essential for period changes following months of T-cycle entrainment (Azzi et al., 2014), this suggests that DNA methylation might universally regulate light-induced period plasticity regardless of the duration of altered lighting paradigms, although DNA methylation might act on different targets for period changes following acute light exposure than those following entrainment. Interestingly, ontological analyses of DNA methylation in the SCN following T-cycle entrainment revealed that the most significant changes in DNA methylation were found in genes encoding neurotransmitter receptors and ion channels (Azzi et al., 2017), suggesting that changes in intercellular coupling within the SCN might be regulated by DNA methylation to induce period plasticity. Further investigating targets of DNA methylation for period plasticity to different lighting conditions will deepen our understanding of epigenetic regulation of circadian clock plasticity.

Lastly, our study has implications in human health. Our data suggest that DNA methylation patterns in the SCN might change dynamically following altered lighting conditions, including a brief light exposure at night or seasonal changes in daylight lengths, to mediate after-effects on circadian period. Given that animals including humans can experience highly varying circadian lighting conditions throughout their lifetime, this suggests that studying dynamics of DNA methylation patterns in the SCN might contribute to understanding the molecular basis of effects of prior light history on the circadian clock (Pittendrigh and Daan, 1976a), and to assessing potential health impacts on human populations experiencing a high degree of changes in environmental lighting conditions including those living at a high latitude with large seasonal changes in daylight and those experiencing circadian misalignments such as shift workers and those frequently traveling across time zones.

## Materials and methods

### Animals and housing

Wild-type C57BL/6J mice (2-4 months old; 000664, Jackson Laboratory) were used for behavioral experiments. For organotypic SCN slice culture from heterozygous PER2::LUC knock-in mice (Yoo et al., 2004), 2-4 months old mice were used in VIP application experiments and P11-14 mice were used in optogenetic stimulation experiments. We used heterozygous PER2::LUC mice as the PER2::LUC knock-in allele can alter circadian functions such as free-running period (Ralph et al., 2021). All animals were housed in a 12:12 light-dark cycle (except as noted), and had food and water provided ad libitum. For circadian behavioral experiments, male mice were used to avoid effects of the estrous cycle on circadian behavior and transferred from their home cages to individually housed wheel-running cages connected to a computer for monitoring locomotor activity using ClockLab software (Actimetrics). Wheel-running cages with ad libitum access to food and water were placed in light-tight chambers and exposed to various lighting conditions controlled by ClockLab including constant darkness and a 16:8 light-dark cycle. For ex vivo assays, both male and female mice were used in experiments. Experiments were performed in accordance with the Vanderbilt University Institutional Animal Care and Use Committee and National Institutes of Health guidelines.

### Stereotaxic surgery and drug delivery to the brain

Mice were anesthetized with 2% isoflurane and placed securely in a stereotaxic apparatus (Kopf Instruments) with the body temperature maintained at 36°C using a homeothermic heating pad (Harvard Apparatus). Eye lubricant was applied to prevent drying during surgery. For RG108 infusion into the third ventricle, an implantable osmotic pump (infusion rate 0.11μl/hr, Model 1004, Alzet) was filled with 200μM RG108 in aCSF (0.66% DMSO) or vehicle and primed with sterile saline at 36.8°C before use as per manufacturer’s instruction. RG108 concentration was chosen based on previous studies examining behavioral outcomes associated with blocking DNA methylation in a brain region (Laplant et al., 2010; Day et al., 2013). An osmotic cannula (28 gauge, Plastics One) was stereotaxically implanted in the third ventricle near the SCN (relative to bregma, A/P: -0.5 mm, M/L: 0.0 mm, D/V: -4.5 mm) and connected to the osmotic pump via a vinyl catheter tubing (Alzet). The osmotic pump was then implanted subcutaneously under the back skin as per manufacturer’s recommendations. After stereotaxic surgery, mice were returned to individually housed wheel-running cages for recovery and subsequent behavioral experiments. To ensure pump infusion, the pump reservoir was inspected after behavioral experiments.

For RG108 injection into the third ventricle, a guide cannula (26 gauge, Plastics One) was implanted in the same location as for infusion, and capped with a dummy cannula (Plastics One). After stereotaxic surgery, mice were returned to individually housed wheel-running cages for recovery and subsequent experiments. On the day of light stimulation in constant darkness, mice were taken out of their cages, and while mice were held tightly under dim red light a dummy cannula was removed and an internal cannula (Plastics One) was inserted into a guide cannula and connected to a 10μl microsyringe (Model 1701, Hamilton) filled with 100μM RG108 in aCSF (0.66% DMSO) or a vehicle via PE50 tubing (Plastics One). Mice were released and allowed to move around while RG108 or a vehicle was dispensed at 0.5μl/sec using a syringe dispenser (Hamilton). After injection, the injection system was disassembled and mice were returned to their cages under dim red light.

### Circadian wheel-running behavioral assays and data analyses

Mice were transferred to individually housed wheel-running cages in light-tight chambers provided with experimental lighting conditions. Mice were habituated in wheel-running cages under a 12:12 LD cycle for at least three days before experiments. For photoperiodic entrainment assays, mice were released into constant darkness for a week to measure a baseline circadian period before re-exposure to a 12:12 LD cycle, and they received stereotaxic brain surgery for RG108 or a vehicle infusion on day 4 or 5 during the 12:12 LD cycle. After additional four or five days in a 12:12 LD cycle, mice were exposed to a 16:8 LD cycle for 12 days and then released into constant darkness to measure an after-effect on circadian period. Light cycle control mice remained in a 12:12 LD cycle for a total of 20 days before a release into constant darkness. For light pulse experiments, mice received stereotaxic brain surgery and after a week of recovery in a 12:12 LD cycle they were released into constant darkness for six days to measure a baseline circadian period. On day 7, mice received RG108 or a vehicle injection under dim red light around CT10.5 (CT12 was defined as the onset of nocturnal locomotor activity) and were given two one-hour light pulses in constant darkness around CT13.5 and CT16.5. Wheel-running activity after the light pulses was recorded to measure phase shifts and period changes.

Wheel-running activity records were extracted and analyzed using ClockLab software running in Matlab (Mathworks). Single- or double-plotted 24-hour actograms with lighting information were produced and activity onsets were determined using ClockLab. Phase shifts were calculated as difference in time between the activity onsets observed following light exposure and those predicted using linear regression from activity onsets before light exposure. Period changes were calculated as differences in the period length of at least six cycles before and after changes in light exposure using Chi-square periodogram in ClockLab.

### Ex vivo SCN bioluminescence rhythm assays

Organotypic SCN slice culture was performed as previously described (Kim and McMahon, 2021). Briefly, 300μm-thick coronal slices containing the SCN were obtained from PER2::LUC mouse brains using a vibratome (Leica) and placed on a semi-permeable membrane insert (PTFE, Millipore) in 35-mm culture dishes. The culture dishes were sealed with a transparent PCR plate film (Bio-Rad) and maintained in a multi-channel luminometer LumiCycle (Actimetrics) inside an incubator at 36.8°C. Bioluminescence from PER2::LUC SCN slices was recorded in 10 min intervals. For VIP application experiments, culture medium was 1.2ml of DMEM supplemented with 10mM HEPES, 25U/ml penicillin/streptomycin, 2% B-27 Plus (Gibco), and 0.1mM D-luciferin sodium salt (Tocris). 1µM VIP (Tocris) dissolved in sterile water was applied at CT14 to the culture medium 15 mins prior to 200µM RG108 (dissolved in 0.66% DMSO) application in the medium. CT12 was defined as the peak of PER2::LUC rhythms. CT14 was determined using at least three cycles of PER2::LUC rhythms before drug application. Drugs were pre-warmed before application and not washed off.

For optogenetic stimulation experiments, culture medium was the same as for VIP application experiments except that it contained 2mM Glutamax (Gibco) instead of L-glutamine. 1µl AAV (pAAV1-Syn-ChrimsonR-tdTomato, Addgene; Klapoetke et al., 2014) was applied onto SCN slices before sealing culture dishes. The opsin expression was confirmed by imaging tdTomato fluorescence 10 days after viral transduction. 10µM SGI-1027 (Tocris) dissolved in DMSO was applied to culture medium one day before optogenetic stimulation. For optogenetic stimulation, 625nm LED light pulses (10Hz, 10ms) were illuminated onto SCN slices at CT14 for 15 minutes using an integrated system of luminometry and optogenetic stimulation previously described (Kim and McMahon, 2021).

### Bioluminescence recording data analysis

Bioluminescence data were analyzed as previously described (Kim and McMahon, 2021). Briefly, raw data were baseline-subtracted and smoothed using LumiCycle Analysis software (Actimetrics). Then they were loaded into Matlab-run ClockLab for further analyses. Phase shifts were calculated as difference in time between the rhythm peaks observed following drug application or optogenetic stimulation and those predicted using linear regression from peaks before manipulation. Period changes were determined as difference in the period length of at least three cycles using linear regression of peaks. Bioluminescence data were visualized using Excel (Microsoft) and Prism (Graphpad).

### Experimental design and statistical analysis

For all experiments, mice were randomly assigned to control and experimental groups. For circadian behavior assays, only male mice were used to avoid estrous cycle effects on circadian period. For ex vivo SCN rhythm assays, both males and females were used. For statistical comparisons, unpaired t-tests, one-way ANOVAs with Tukey’s post hoc tests, or two-way ANOVAs with Sidak’s post hoc tests were performed using Prism, and tests used for individual experiments are described in the figure legends. Data are presented as mean ± standard error of mean (SEM). Differences between groups were considered statistically significant when the p-value was less than 0.05.

